# Y chromosome STR variation reveals traditional occupation based population structure in India

**DOI:** 10.1101/2024.08.28.610024

**Authors:** Jaison Jeevan Sequeira, M. Chaitra, Ananya Rai N R, M. Sudeepthi, R. Shalini, Mohammed S Mustak, Jagriti Khanna, Shivkant Sharma, Rajendra V E Chilukuri, George van Driem, Pankaj Shrivastava

**Author notes:** Correspondence: Jaison Jeevan Sequeira. George van Driem.

## Abstract

Earlier models of grouping Indian populations were based on language families, social stratification and geographical location. Such grouping system has often resulted in oversimplification of ancestry inferences. Moreover, we do not find many studies focused on studying the variation within these groups and the role of past demographic events in shaping them. We analysed the Y-chromosome Short Tandem Repeats haplotypes from 8153 males from India and Eurasia to explore the impact of Holocene migration on the Indian gene pool. We used haplotype variation and date estimates to understand the characteristics of each haplogroup with respect to the different grouping models. Our findings show that the Neolithic agricultural expansion has had a strong influence in shaping the male gene pool of the Indian subcontinent. Haplogroups F, L and R1a contribute greatly towards stratifying Indian populations as hunter-gatherer related, farming-related and priestly groups respectively. Although the caste system enforced endogamy, a traditional occupation based admixture existed since the Neolithic times. Dispersal of haplogroup L from the Near East played a major role in the formation of an agriculturist population that formed an intermediary between the primitive tribes and the R1a-rich priestly group. This study shows that the frequency of R1a in the hunter-gatherer tribes (1.5%) is much lower than previously reported based on other models of population clustering.

## Introduction

The modern-day Indian gene pool derives its ancestral components from the Andamanese hunter-gatherer related ancestry, farmer-related ancestry from the Iranian plateau and the Pontic-Steppe pastoralist-related ancestry ^1,2^. However, reality is much more complex with many temporal layers. Studies have shown ancestral components unique to different cultures and geographical regions at different time depths ^3–6^. Although these models lack concrete support from the archaeological record, the genetic components found in the Indus Valley, Swat Valley, Bactria-Margiana samples do not fit perfectly within the existing models of ancestral reconstruction used for the Indian subcontinent ^3,4^. A recent study identified a Proto-Dravidian ancestral component in the Koraga tribe from the southwestern coast of India ^7^. The proxies used for ghost population components like Ancient Ancestral South Indian (AASI) need validation from ancient DNA data. The existing models use Andamanese genomes as a proxy for the hunter-gatherer ancestry in India. The substructure within this widespread ancestry and the impact of genetic drift in the Andamanese islanders have been largely ignored. Ancestral reconstruction studies based on a few samples from the Indian subcontinent have left many questions unanswered and have moreover resulted in oversimplification.

One of the best examples is the case of haplogroup R1a. This haplogroup is prevalent in the Indian subcontinent, particularly among Indo-European speakers, and even South Indian tribes exhibit a moderately higher frequency of R1a ^8,9^. The presence of R1a, a haplogroup often linked to Indo-Aryan languages and the Aryan Migration Theory, in primitive Dravidian-speaking tribes dovetails with the reconstruction of the complex social history of the Indian subcontinent arrived at by linguists ^10^. Previous studies have suggested Central Asia and West Asia as potential points of origin for the R1a variant ^11,12^, while others have proposed its origin in India ^8^. A seminal paper in 2009 tried to cluster the modern-day Indians into Ancestral North Indians and Ancestral South Indians depending on the gradient of West Eurasian component and Andamanese hunter-gatherer component respectively ^1^. Since then, studies have attempted to correlate the admixture pattern with caste based endogamy that was introduced about 1,900 and 4,200 years ago ^13^ and others have argued for the existence of such endogamy since the Neolithic times ^14^. Such ambiguities exist even today due to the lack of ancient genomes from these time periods. Is it reasonable to assume that waves of Indo-European males carrying the R1a haplotype replaced the male gene pool in the Indian subcontinent? Studies, alternatively, have also identified tribes without any West Eurasian haplogroups in the same geographical area inhabited by tribes with moderate levels of R1a ^14–16^. Is this inconsistency due to inadequate clustering models?

Another important demographic event that impacted the gene pool of Eurasia and Africa is the Neolithic agricultural expansion ^17,18^. Although about a half of the Indian gene pool is derived from the farmer-related Neolithic component ^2^, the impact of this event has not been studied enough. Recent studies have identified the Near East as the population hub about 40kya ^19,20^. Archaeological and linguistic records document the ancient civilisations in this part of the world. Ancient DNA data emerging from the Iranian plateau provide a suitable proxy for the early Neolithic farmer gene pool ^21,22^. Due to the availability of ancient DNA in Europe, recent studies were able to reconstruct the impact of agricultural expansion in Anatolia and Europe ^21,23,24^. However, the case is not so promising in the Indian subcontinent due to a lack of good-quality fossils. The only prospect appears to lie in the endogamy of modern-day Indian populations that may have conserved chromosomal traits from their Neolithic ancestors. To study them, a huge number of whole genomes must be sequenced, which presents us with today’s challenge. Revisiting existing data from published autosomal and uniparental chromosomes is another options left to answer these pertinent questions of the past.

In this study, we examine the haplotype variation within each haplogroup and compare this variation with different models of sample stratification. For this purpose, we use the Y-STR allelic data from 8153 males from India, Iran and Eurasia. Short Tandem Repeats (STRs) have been used extensively for forensic applications, and in recent years the number of loci included in forensic kits have been increasing. Currently, Y-STR kits use at least 23 markers, including 15 common markers in the old and new kits. In addition to forensic applications, these markers carry ancestry information as well. Due to their faster mutation rate and non-recombining nature, Y-STRs provide a reliable overview of the past. Early human migrations were initially gender-neutral, but during the Holocene, there was a shift towards male-dominated migrations ^25,26^. Was there a complete overhaul in the male gene pool in the Bronze age? What was the impact of agricultural expansion on the primitive tribes that followed pre-Neolithic occupation? In the present study, we will attempt to provide a more resolved picture of the haplotype variation in the Indian subcontinent using Y-STR data. We will utilise a population modeling approach based on traditional occupations to characterise Indian populations ^14^ and compare this approach against existing models based on linguistic affiliation, social hierarchy and geographical location.

## Materials and methods Sample information

Y-STR data from 8153 males from India, Iran and Eurasia were pooled from earlier studies ^14,27,36–43,28–35^ (ESM2 Table S6 and S7). To compensate for the lack of data from the south-west coast of India, we genotyped 81 males (ESM2 Table S5). 3 ml blood samples were collected from healthy, unrelated individuals with a written consent. The study was approved by the Institutional Ethics Committee at Mangalore University (MU-IHEC-2020-3 dated 21.08.2020). All methods were performed in accordance with the Declaration of Helsinki and regulations of the Ethical Committee. We classified the Indian samples into three models. The model based on social hierarchy operates on the basis of the two broad groups of castes and tribes. The model based on linguistic affiliation groups samples into Indo-European and Dravidian. We have selected only these two language families due to their relevance to our objective. The occupation-based model groups samples based on the traditional occupations with which they are known to be associated. This approach was used by Arunkumar et al. 2012 to study the male lineages in the south Indian castes and tribes ^14^. The hunter-gatherer-related populations follow pre-Neolithic practices like hunting, gathering and fishing. The populations involved in dryland farming are considered under farming-related group. The Brahmin communities, who are traditionally associated with religious rituals, are categorised as priestly group. These groups are further clustered under geographical regions if populations following the same traditional occupation are separated by a large distance.

### Genotyping

DNA was extracted using standard extraction procedure described elsewhere. PCR reagents were standardized and amplification was performed using a 9700-thermal cycler (Applied Biosystems, USA). The PCR mix was prepared using Ingenomics™ Y Profiler 2x Master Mix V- 6.25μl, Ingenomics™ Y Profiler 5x Primer Mix- 2.5 μl, and MQ water 3.75 μl for one 1.2 mm washed punch of FTA paper. Genotyping was performed according to the manufacturer’s protocol included with the kit. In brief, the PCR products were genotyped using capillary electrophoresis with POP-4 polymer in an ABI 3500 Genetic Analyzer (Applied Biosystem, Foster City, CA, USA). GeneMapper ID-X Software v1.6 was used to analyse the data to identify alleles through comparison with the allelic ladder included in the kit. For allele designation, the peak detection threshold was chosen at 100 RFUs (Applied Biosystem, Foster City, CA, USA). To assure a high level of quality, a positive control and a negative control were run along with every batch of samples.

### Statistical analysis

#### PCA based on haplogroup frequency distribution

Haplogroups were determined for all the samples using Athey’s Y-haplogroup predictor ^44^. Principal component analysis was performed using *prcomp* function in R ^45^. *ggbiplot* package was used for visualisation in R.

#### RST based Multidimensional Scaling (MDS) and Heatmap

Reynold’s pairwise distance (RST) was calculated for each population using the *adegenet* package in R for RST based Multidimensional Scaling and Heatmap ^46,47^. The distance matrix was visualised using the *pheatmap* package in R.

#### Haplotype network analysis

We used NETWORK 10.2.0.0 software (http://www.fluxusengineering) to build a haplotype-based network ^48^. We entered the allele count values manually into the software and ran the median joining algorithm ^49^. Samples were pre-processed using the star-contraction algorithm in default settings. Post processing was done using the MP option to purge superfluous links and median vectors not contained in the shortest trees in the network. The nodes were coloured as per the traditional occupation categories. Samples from Europe and Iran were also coloured to understand the lineage sharing pattern.

#### Estimation of TMRCA using weighted rho (**ρ**) statistic

The estimation of the time to the most recent common ancestor (TMRCA) was based on Y-STR haplotypes using a protocol described elsewhere ^50,51^. Haplogroups were dated, based on STR variation with ρW, a weighted version of ρ that leverages the relatively precise knowledge of the mutation rate of each Y-STR. The weighted rho (ρW) were calculated using an R script available on GitHub (http://github.com/fcalafell/weighted_rho). Mutation rates were obtained from the Y-Chromosome STR Haplotype Database. All analyses were performed using only complete haplotypes (all 15 loci namely DYS19, DYS389I, DYS389II, DYS390, DYS391, DYS392, DYS393, DYS437, DYS438, DYS439, DYS448, DYS456, DYS458, DYS635, GATA H4). DYS385 was omitted from the analysis due to the inability to differentiate between the DYS385a and DYS385b loci with the Y STR kit. The number of repeats at DYS389II was derived by subtracting the number of repeats at DYS389I. Treemap was plotted using MS Excel ^52^.

#### Y-SNP based analysis

Haplogroup distribution data were obtained from 59 populations (n=3605) from South Asia, grouped by their language family, geographical location and traditional occupation. PCA was performed using the prcomp function in R software. AMOVA was performed using Arlequin software ^53^. QGIS was used to generate contour maps ^54^.

### Results and discussion

#### Population structure inferred from clustering models

We pooled a dataset of 8153 males from India, Iran and Eurasia (Table 1). For the Indian samples, we collected information about their social affiliation, linguistic classification and traditional occupation (ESM1 Table S1). Mixed samples without this information were excluded from specific analyses.

**Table 1.** Sample information

|  |  | No. of Samples | No. of populations |
| --- | --- | --- | --- |
| **Indian** |  | 5187 | 50 |
| Social | Castes | 1992 | 26 |
|  | Tribes | 1490 | 16 |
| Linguistic | Indo-European | 1597 | 12 |
|  | Dravidian | 2533 | 34 |
| Occupation-based | Hunter-gatherer | 1490 | 16 |
|  | Farming-related | 733 | 9 |
|  | Priestly | 578 | 8 |
| **Global** |  | 2966 |  |
| Iran | Iran West | 577 |  |
|  | Iran East | 256 |  |
|  | Iran North | 415 |  |
|  | Iran South | 105 |  |
| Europe |  | 325 |  |
| Central Asia |  | 282 |  |

From the principal component analysis, it is evident that the studied populations show resolved structure when stratified by traditional occupation (Fig. 1). The major contributors to this variation are haplogroup F, R1a and L, which correspond with hunter-gatherers, priestly communities and farming groups respectively. Earlier studies qualified haplogroup F and its descendant haplogroup H as ‘Indian-specific’ ^9,55^. This observation is due to higher frequencies of these haplogroups in the tribal populations. Similarly, haplogroup J is widely accepted as a marker for the agricultural expansion into India ^17,56^. In our study, we find that the spread of agriculture from the Iranian plateau is strongly associated with haplogroup L. This is in congruence with the findings of a recent study on Y-chromosomes from South Asia linking L-M22 marker with the spread of agriculture as well as with the Dravidian languages into the Indian peninsula ^57^. Haplogroup R1a on the other hand, is often associated with the spread of Indo-European languages ^58,59^. Ancient DNA studies show that this male lineage started spreading from the Pontic-Caspian region about 5,000 years ago ^60^ both westward and eastward, reaching the Indian subcontinent about 3,000 years ago ^3^. A recent study involving large-scale whole-genome data has identified three major ancestral components in Indians namely, Andamanese hunter-gatherer related ancestry, farmer-related ancestry from the Iranian plateau and Pontic-Steppe related ancestry ^2,3^. These sources correspond with our PCA result showing maximum structure when samples are grouped by traditional occupation. Overall, it is evident that the modern-day population clusters formed mainly through male-biased admixture.

**Fig. 1.**
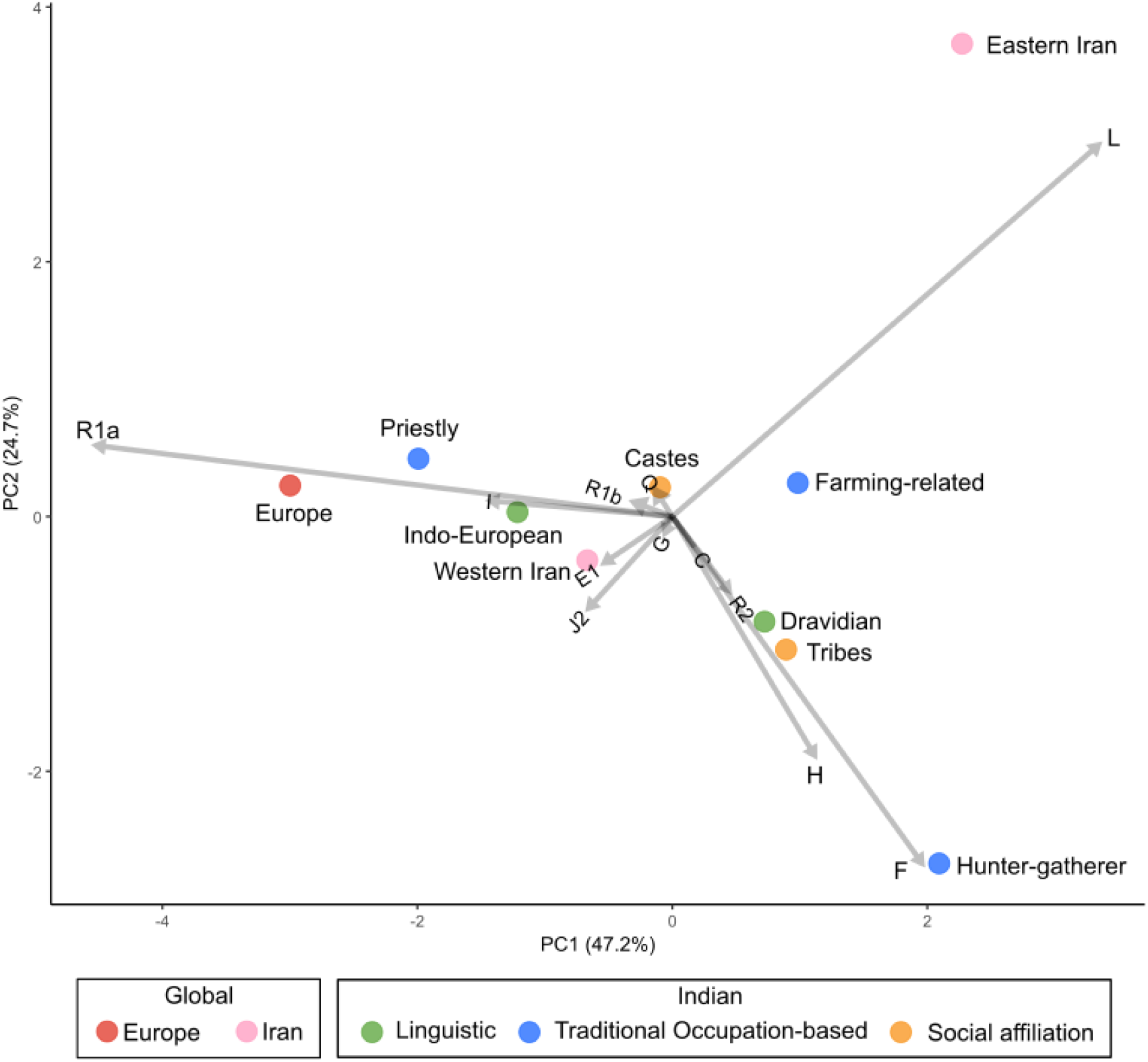
PCA plot showing population clusters based on Y-haplogroup frequency distribution in the India, Europe and Iran. Biplot represents variance contribution of haplogroups. Coordinates in the biplot are coloured based on linguistic affiliation, traditional occupation and social stratification in the Indian populations.

#### Haplotype landscape within India

To explore the variation in individuals within each haplogroup, we plotted an RST-based Heatmap (Fig. 2). Broadly, three major clades were observed within the R1a haplogroup. The R1a haplotype from Eastern Iran and Kyrgyzstan formed a separate cluster showing high pairwise distance between themselves and the rest of the R1a haplotypes. The second cluster is formed by the European and Altaian and Kakhassian R1a haplotypes. Interestingly, Western Brahmins and Caucasians show affinity towards the Altian and Khakassian samples. We have reported a similar case in the maternal ancestry of Citrapur Sārasvat Brahmins in their N macrohaplogroup lineage (Sequeira et al. 2024 unpublished). The third cluster includes all other R1a haplotypes from India, Afghanistan, Caucasus and Iran. Whilst the farming-related and priestly groups cluster together, the R1a haplotypes in the hunter-gatherer groups cluster away from them. This observation leads us to suggest that paternal lineage sharing was more frequent among groups of similar social status. The wide gap between the frequency of R1a in the hunter-gatherer tribes (1.5%) and the priestly group (43.4%) (Fig. 3) further substantiates our claim. Such a contrast is not observed when these populations are clustered based on social status (castes and tribes) or linguistic affiliation (Indo-European and Dravidian) ^8,9^ prompting the authors to advance hypotheses such as widespread admixture following the Indo-Aryan migration or an Out-of-India migration. Ambiguous grouping strategies led to diverse patterns of haplogroup frequency, showing that the genetic isolation of ancient tribes was compromised. Strong founder intensity and inbreeding reported in many tribal groups of India ^61,62^ support our hypothesis. The role of agricultural expansion in both the assimilation and displacement of pre-existing tribes in South India ^14^ deserves more attention.

**Fig. 2.**
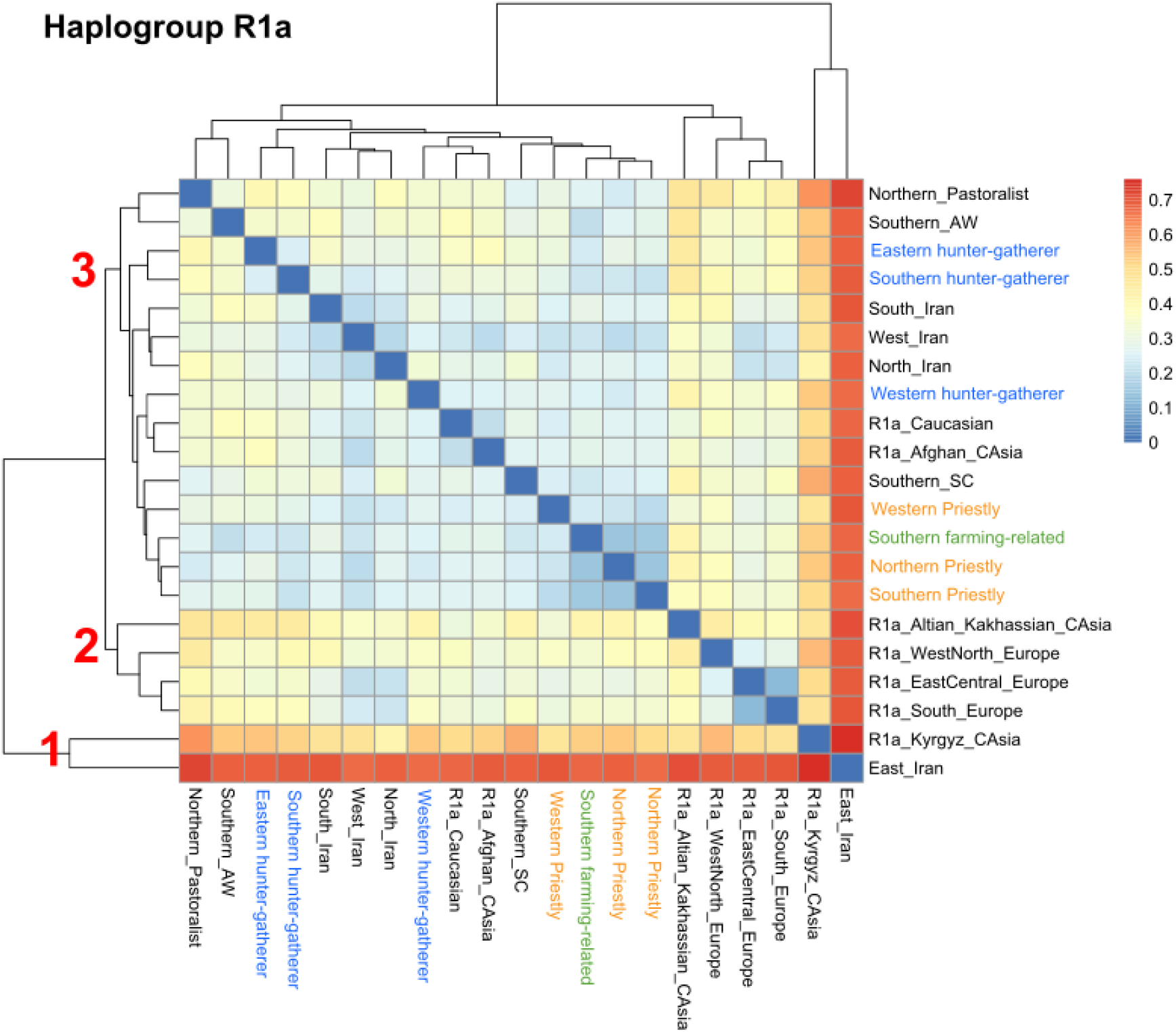
RST based Heatmap for R1a samples. Three clusters are shown in the figure. Blue represents hunter-gatherer group, orange represents priestly group and green represents farming-related group.

**Fig. 3.**
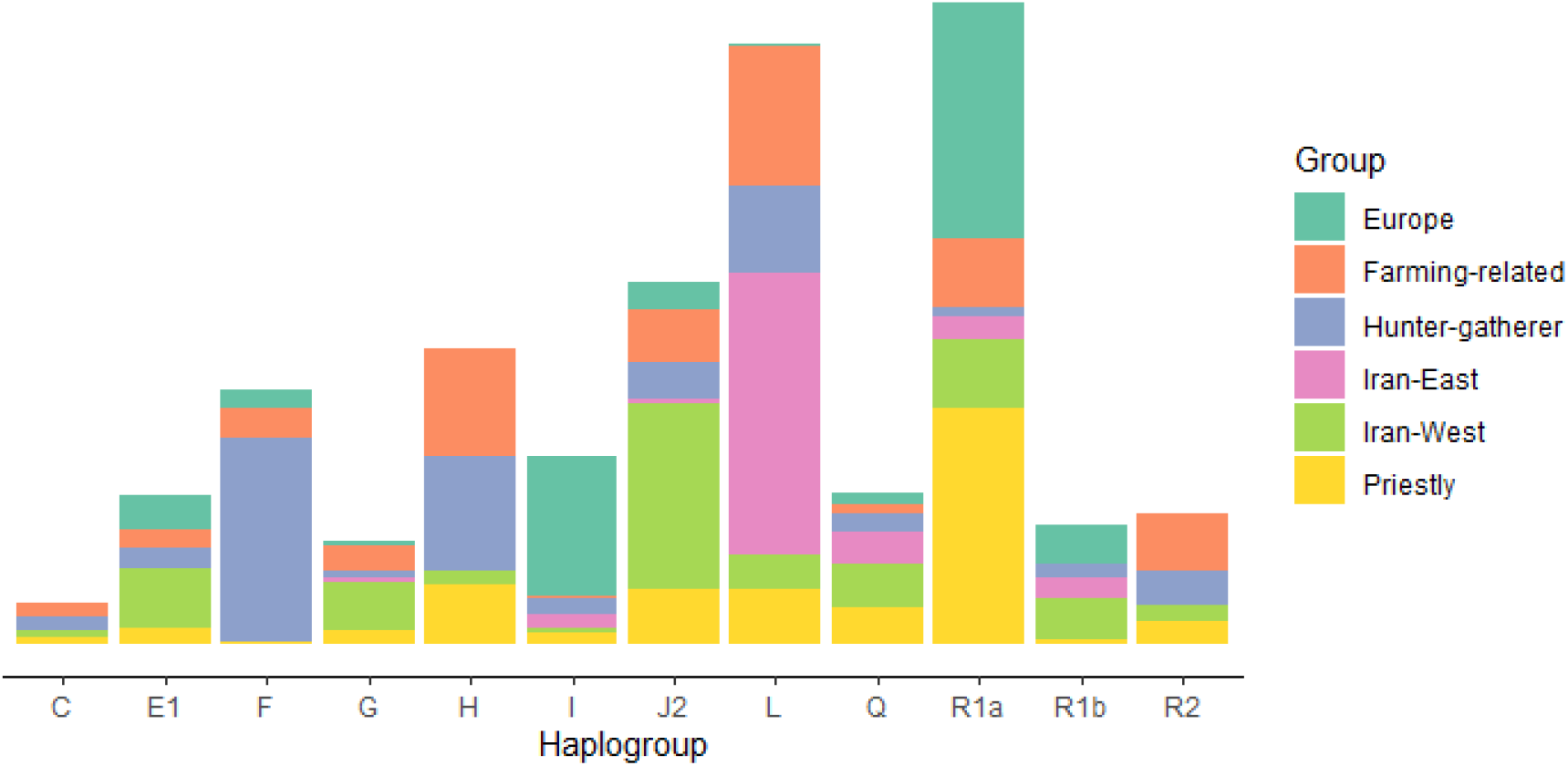
Stacked bar plot showing haplogroup distribution pattern in the studied groups

The gradient in the frequency of haplogroup F provides additional evidence for population structure based on traditional occupation. Haplogroup F levels peak at 37.7% in the hunter-gatherer tribes (Fig. 3). Haplogroup F accounts for 5.5% of paternal lineages in farming-related groups. Haplogroup F is almost absent in the priestly group (0.2%). However, its descendent haplogroup H is distributed among all the three groups at frequencies between 10-21%. The RST-based Heatmap shows only two major clusters for haplogroup H (EMS1 Figure S1). We observe that the farming-related group and the hunter-gatherer group share the same clade, whilst the priestly groups cluster away. H haplotypes found in eastern and western hunter-gatherer tribes are distinct from those found in southern hunter-gatherer tribes. Among F and H haplogroups, only F seems to be localised in South India. Haplogroup H, on the other hand, has a wider presence, stretching between the Iranian plateau and the Indian peninsula. Haplogroup H is also reported in Bronze Age individuals from present-day Iran and Turkmenistan ^3^. Earlier studies qualified haplogroup H as India-specific ^56^. The longitudinal correlation with geography observed for haplogroup H ^56^ and its abundance in the western part of the Indian subcontinent suggests that haplogroup H may have been the most common lineage that flourished in the region between the Iranian plateau and the Indian peninsula after splitting off of haplogroup F, which itself represents a remnant of the earliest migration along the Southern Coastal Route ^16^.

We observe three major clusters within haplogroup L (EMS1 Figure S1). Cluster 1, which includes Eastern Iranian population and hunter-gatherer groups from western and eastern India, forms a distinct clade due to dissimilarity with rest of the L haplotypes. Cluster 2 includes populations from southern India, whilst Cluster 3 includes populations from Iran and North India. The farming-related group and the southern hunter-gatherer tribes belong to the same clade, whereas the northern priestly group clustered with the pastoralist Gujjar population and Iranian groups. From this, it is evident that all the Indian populations bearing the L lineage originated in West Asia ^57^ and spread into the Indian subcontinent during the Neolithic expansion.

Haplogroups R2 and J2 are found in farming-related groups at a frequency of 9-10% (Fig. 3). The J2 haplogroup which peaks in western Iran is distributed evenly in the farming-related and priestly communities at around 10%. The presence of haplogroups L, J2 and R2 in the priestly and farming related groups and the contrast in the frequency of R1a in these groups (43.4% vs. 13%) suggest that the basal component of the male gene pool of most present day non-tribal Indian groups is related to Neolithic migrations from the Iranian plateau. The L lineage diverged into a Near Eastern cluster and an Indian cluster which further diverged into northern and southern lineages.

#### Inference from the TMRCA of haplotypes

The haplotypes found in the predefined groups were dated using a weighted rho statistic described elsewhere ^50^ (ESM2 Table S8). The oldest TMRCA in the Iranian haplotype dates back to the time of the Last Glacial Maximum (LGM) (Fig. 4). However, we observe a contrast in the age of haplotype in the samples from Eastern and Western Iran with the former showing deeper lineages than the latter.

**Fig. 4.**
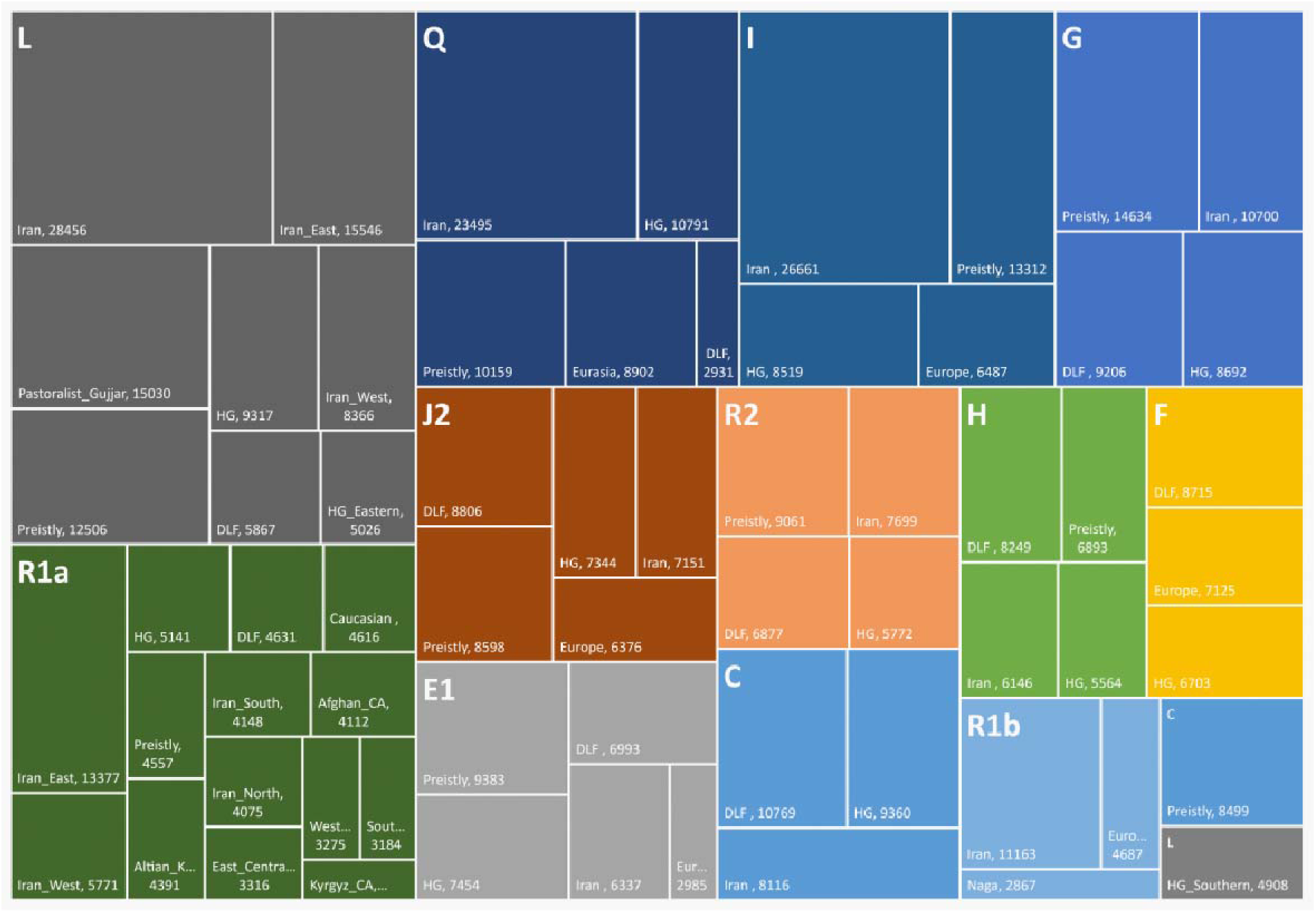
Treemap showing TMRCA of haplogroups found in the studied population groups

The oldest ancestral lineage of haplogroup R1a is found in Eastern Iran (∼13,000 ybp). We observe a difference of about 7,000 years in the TMRCA of Eastern and Western Iranian individuals, the latter being younger. The age of R1a halplotype in Northern Iranian and Caucasian individuals dates from about 4,000 years ago, suggesting a clinal connection that may have involved the formation of the Grave Pit culture male gene pool ^60^. In Indian groups, the R1a haplotype is about 4,000 to 6,000 years old. The R1a lineage found in the priestly group (4557 ybp) is the youngest in India. However, the TMRCA of R1a found in Indian groups is more similar in age to the haplotype found in Iran (North, South, and West), the Altai, Caucasus and Afghanistan as compared to the Eastern Iranian haplotype. The impact of East Eurasians and Central Asians is known to have been greater on the Eastern Iranian frontier, which may explain the presence of older haplotypes in this region ^63,64^. On the other hand, the R1a lineage found in Europe and Central Asia is much younger (2000-3300 ybp), which may be due to the spread of cultures descending from the Grave Pit cultural complex in these regions ^3^, completing the loop of haplogroup R which originated in Central Asia ^12^. Our findings indicate spatio- temporal differences in the age of R1a, with older lineages in Iran and India, and younger lineages in Europe.

Interestingly, we do not observe any haplotypes among the Indian groups that date back to the LGM or pre-LGM era. This finding contrasts with the age of maternal haplogroups found in India ^65^. Haplotypes H, F, C and J2 found in the farming-related groups of India are about 8,000 to 10,000 years old. A similar age for these haplotypes is found in hunter-gatherer tribes. Haplotype L in the Gujjar pastoralist group is about 15,000 years old. The TMRCA of the same haplogroup in the priestly communities is ∼12,500 ybp. The oldest lineages of Q, I, G, R2 and E1 are also found in the priestly group. The presence of these haplogroups, with plausible origin in Europe and Iran, in the priestly communities suggests lineage sharing between the ancestors of present-day priestly groups and the West Eurasians at a deeper time depth. Also, the high frequency of the younger R1a lineage (∼4500 years old) in the priestly group coincides with the timeline of the Aryan influx, which in turn explains the presence of the relatively higher proportion of Pontic-Steppe-related ancestry in the northern Indo-European caste groups ^2^. The presence of all the other older lineages of West Eurasian haplogroups is in congruence with the reported moderate levels of Neolithic ancestry in the Indo- European speaking northern populations ^2^, which also characterises the priestly group studied here. It is also important to note the difference between the Western priestly group, which includes Sārasvata Brahmins, and other Northern and Southern priestly groups. The RST-based Heatmap shows greater affinity between the R1a haplotype found in the Western priestly group and the Altaian population. Whilst this affinity is interesting, a whole-genome study may provide a more resolved picture of the fine-structure variation that exists within the priestly communities.

With the exception of haplogroups R1a, L and F, which contribute most to the variation in the traditional occupation based groups, all other haplogroups are ubiquitously present in all Indian castes and tribes. This distribution may be due to admixture before the introduction of strict endogamy in India. However, even before the introduction of caste system, admixture between these groups appears to have been more frequent as compared to the priestly communities based on the similarity observed in the TMRCA of haplotypes of farming-related groups and hunter-gatherer tribes. We propose that about 12,000 years ago, some hunter-gatherer tribes in the Iranian plateau evolved into farming groups forming the sedentary Neolithic civilisation. This transition resulted in the assimilation of haplogroup L which dispersed towards the Deccan region in peninsular India. These early farmers mixed with the H haplogroup carrying tribes and then formed the ASI component. The ASI farmers heralded the South Indian Neolithic. The farming group localised in the Indus valley diversified after diverging from the Early Neolithic farmers through admixture (assimilating older R1a, G, R2, E, H and J) and formed the Harappan civilisation (Indus cline). The pastoralists from the north (the Pit Grave culture descendants with R1b, I, younger R1a and Q) admixed with the farming- related Indus people and formed the ANI component. This group introduced kingship and maintained endogamy. A group diverged from this ANI cluster and formed the priestly class and maintained endogamy, resulting in a retention of >40% R1a. The F haplogroup carrying tribes remained relatively isolated, retaining the AASI component.

#### Lineage sharing network within the occupation-based groups

To understand the lineage sharing pattern within each of the haplogroups, we constructed haplotype networks for major haplogroups found in the studied population (Fig. 5). The R1a network shows a branch specific to the European haplotypes. The branch arises from a haplotype shared by Iranians and the priestly group. This finding further substantiates the affinity of the priestly group with the European samples. However, the model suggests more than just a Bronze Age admixture. Since the basal haplotypes for R1a found in all other groups are from the Iranian plateau, we may infer that R1a spread into Europe and the Indian subcontinent in the first instance (spreading the older R1a) followed by another instance of assimilation due to endogamy. Consequently, we observe Iranian and hunter- gatherer males sharing basal haplotypes with the priestly group. The presence of basal haplotypes in Iran is also seen in haplogroups E, J, Q and L. In the J and L haplogroup networks, we find Iran- specific clusters suggesting events of assimilation. The trend in haplogroup J is similar to that of haplogroup R1a and E, where we find basal haplotypes in the Iranian males. In case of J, basal haplotypes in the farming-related group are found as well. A separate clade of L (probably L-M76) seems to have formed in the farming-related group ^57^. In addition, we find basal haplotypes of L arising in the hunter-gatherer group. In case of priestly group, we observe basal haplotypes in haplogroup Q, R1a (shared with hunter-gatherer), R2, H and L. Eventually, similar haplotypes assimilated in the priestly group probably due to strict endogamy. This finding suggests that the male gene pool in the present-day priestly group preserves remnants of Neolithic ancestry along with the European Bronze Age source, as is evident from the lineage sharing observed on the tips of the R1a tree. Our R1a network also shows that the Bronze Age admixture was not universal. This haplogroup was already present in the Iranian plateau when a more diverse Neolithic culture developed in this region. Haplogroup F exclusively originated in the hunter-gatherer group and shows at least three different clusters. Haplogroup H on the other hand shows basal haplotypes in farming-related and priestly groups. We observed a lot of haplotype assimilation in the Gujjars, who are traditionally pastoralists. We have not included pastoralists in our model to avoid ambiguities that may arise in considering the farming related ancestral population that migrated more slowly as compared to the nomadic herding tribes ^10^. This hypothetical scenario requires further investigation.

**Fig. 5.**
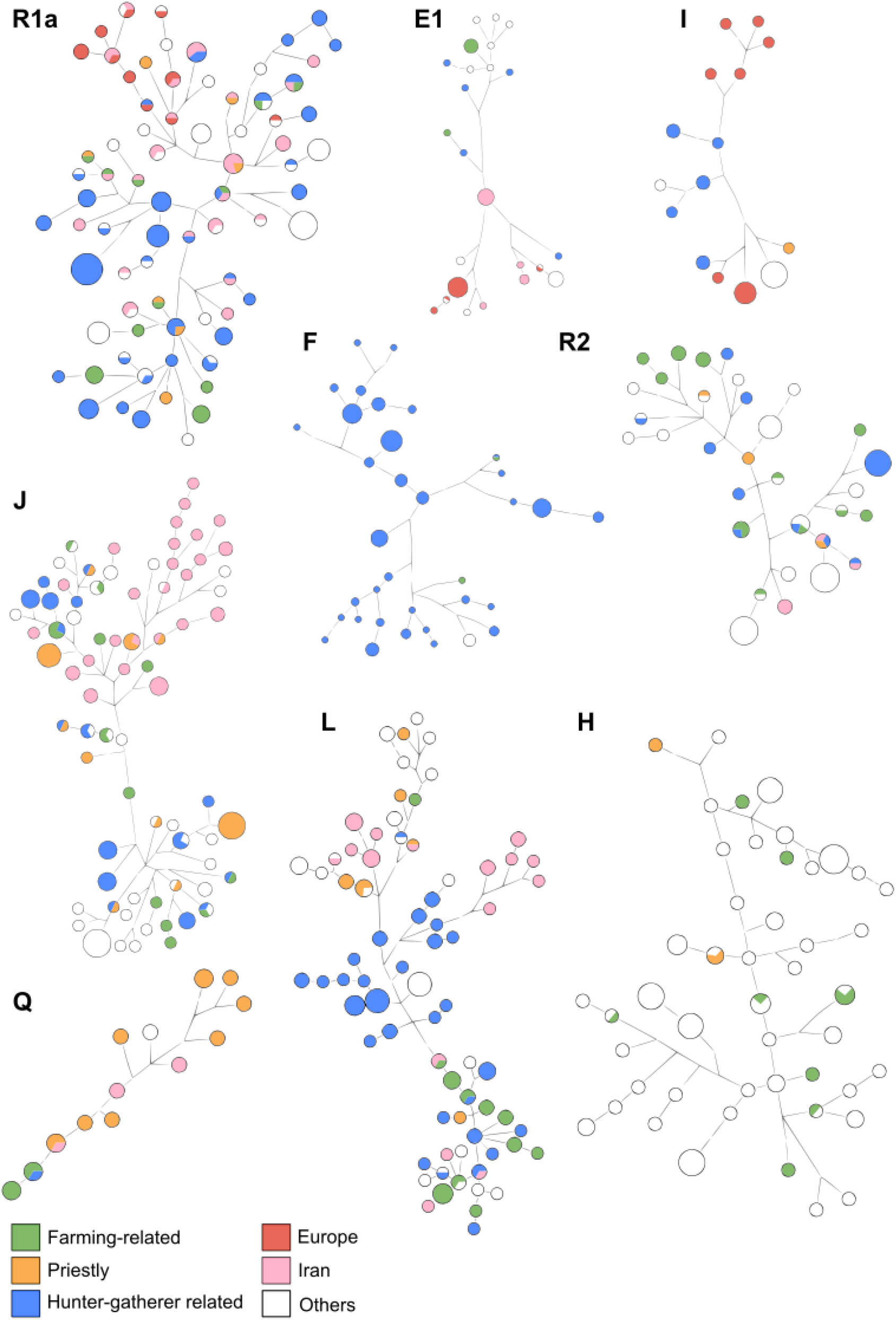
Haplotype network in major haplogroups

#### Association of haplogroup L with the Neolithic agricultural expansion

Haplogroup L-M20 is the descendant of K-M9, which originated closer to the Middle East 50,000 years ago ^66^. The time estimate for the L-M20 subclade in Pakistan is ∼26,000 years ^67^. Whereas its age estimate in the South Indian agriculturist Okkaliga and Li□gāyata population is between 15,000 to 18,000 years ^68^. A similar coalescent age is reported for L-M20 found in Afghanistan. Therefore, L- M20 possibly originated in Pakistan, spread into India during the Last Glacial Maximum (LGM) recovery period and formed L-M27/M76, which today is found mostly in Dravidian tribes and castes directly or indirectly involved in agriculture at frequency averaging around 22% (ESM1 Figure S1 and ESM2 Table S1). L-M27 found in castes and tribes of Tamil Nadu is 15,000 years old. Interestingly, the hill tribes found in the same region belong to a different gene pool, with pre-LGM haplogroups in their maternal as well as paternal lineages ^14,16,65^. Moreover, the caste groups that live in this region as scheduled castes, mixed with these tribes and exhibit a lower R1a frequency (<20%) and moderate levels of H, J and L. This pattern suggests that paternal lineage sharing was common in the lower strata, whereas such paternal lineage sharing has been uncommon between the priestly groups and the hill tribes at least since 7,000 years ago ^14^. Our findings show that the agricultural communities occupy an intermediate position between the hunter-gatherer related and priestly groups. Therefore, we suggest that the Neolithic agriculture linked with haplogroup L, probably related to the early agriculturists who first inhabited the Iranian plateau, resulted in a social hierarchy in South India which was anterior to the social arrangement born out of Bronze Age admixture.

When we model these populations by linguistic affiliation, this fine structure variation gets veiled. This is because all the South Indian groups speak a Dravidian language irrespective of their traditional occupation or social status. We could quantify the potential of population stratification by traditional occupation by comparing the AMOVA results (ESM1 Figure S3 and ESM2 Table S2). AMOVA showed similar percentage variation when populations were grouped by traditional occupation, linguistic affiliation and geographical location. However, population differentiation was slightly lower among groups (FCT=0.039) that followed a similar traditional occupation. This pattern may be due to higher diversity in these populations. Nevertheless, the influence of Neolithic agriculture on the gene pool of present-day Dravidians is evdiently similar to that of recent admixture with R1a component, which is relatively higher among Indo-European populations and the Dravidian priestly communities.

Occupation-based stratification is also observed in Y-SNP multivariate analyses (Fig. 6). Data were broadly classified by traditional occupation. Haplogroups that were earlier associated with the Indian subcontinent, West Eurasia and Neolithic agriculture were pooled for multivariate analysis. We observe a V-shaped distribution with the hunter-gather-related and priestly groups occupying the extremities. The farming-related groups cluster at the converging region with greater overlap with the non-farming cluster. In order to understand which haplogroups are responsible for this clustering pattern, we applied gradient colouring based on the frequency of different haplogroups (S4). We made four groups considering the existing information of their origin, divergence time and association with demographic events. Haplogroups F, H, R2 and C5 predate the Neolithic period in India. R1a is associated with the spread of Indo-European languages and the Bronze Age migration. Haplogroup J is believed to have dispersed with the spread of agriculture and was grouped with G and L3-M357 due to their geographic proximity. All these three groups showed different patterns of gradient on the PCA plot. Interestingly, the pattern observed for haplogroup L-M27/76 is very similar to the pattern observed except that the overlap of R1a with L is less than with J.

**Fig. 6.**
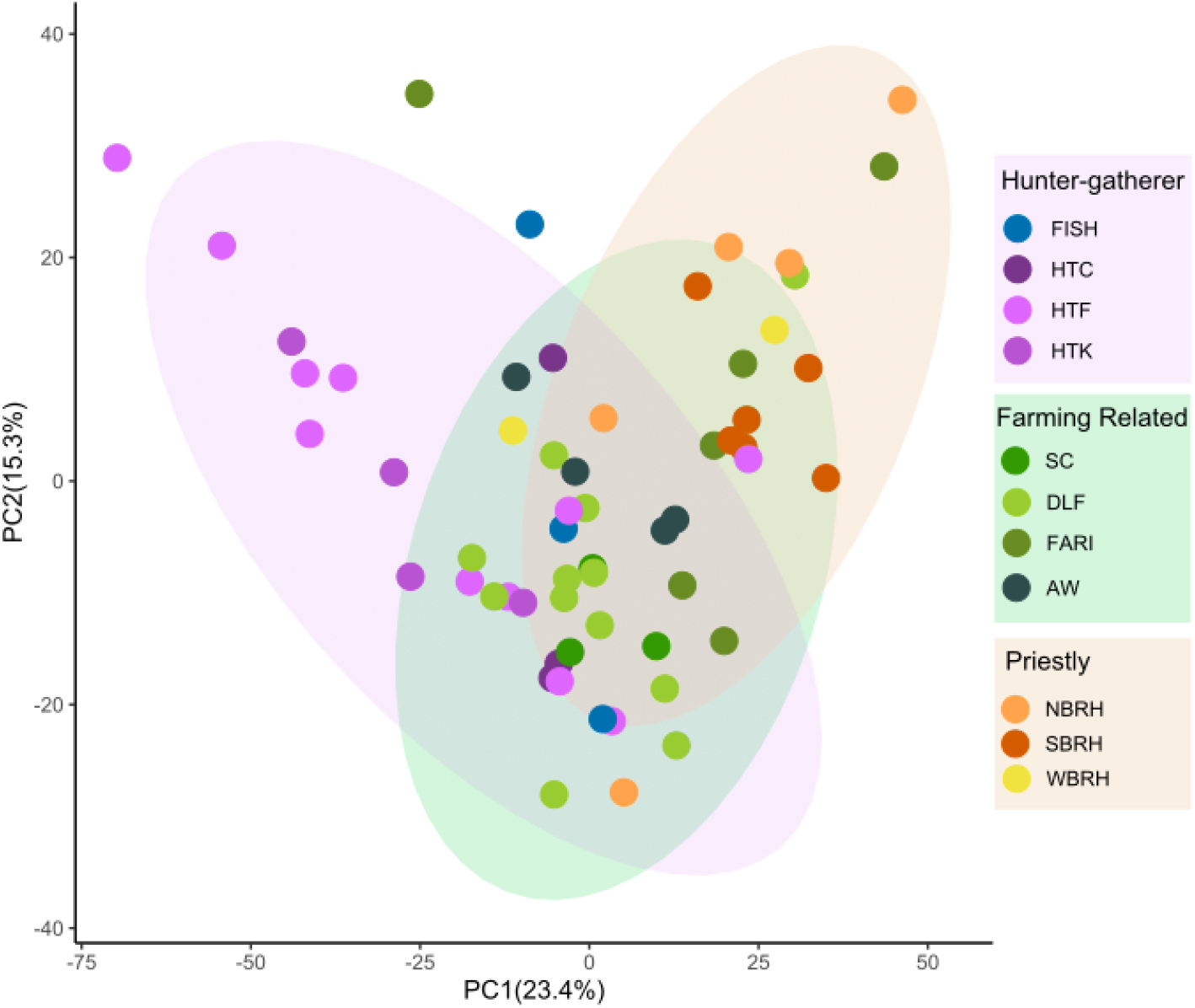
PCA plot showing clustering pattern based on traditional occupation. FISH: Fishing communities, HTC*, HTF* and HTK*: Hunter-gatherer tribes, SC: Scheduled Castes, DLF*: Dryland Farmers, FARI: Paddy cultivating farmers, AW*: Artisans and Warriors, NBRH: Northern Brahmins, SBRH: Southern Brahmins, WBRH: Western Brahmins. *These codes are same as Arunkumar et al. 2012.

Although the time depth of early pastoralism and ash mound culture in the Southern Neolithic is estimated to date from 3,000 BC, sheep domestication in the Mehrgarh Period I (dated to earlier than 7,000 BC) and the cultural continuity between the northwest and the Chalcolithic North Deccan suggests that the agriculturalists/pastoralists of the south separated from the Indus cline corresponding with the bifurcation of L-M20 into L-M27/M76 (∼10,000 years ago). Based on the mtDNA haplogroup distribution pre and post LGM, we also propose that diversification in the gene pool of the Indian subcontinent is heavily influenced by early Neolithic agriculture. Therefore, civilisational developments in and around the Indus Valley started much earlier than the present estimate and may be associated with an earlier perhaps proto-Elamo-Dravidian agriculturalist expansion. As we prepare to interpret vast amounts of DNA data from the Indian subcontinent, diverse factors and grouping strategies must be considered. Informed sampling methods will prove very useful in avoiding sampling errors at the initial stage of data generation. Our findings show that the traditional occupation is an important factor to consider when studying the ancestry of the Indian populations. L- M27/M76, an Indian-specific subclade of L-M20, linked with Neolithic agriculture, is therefore of great relevance in this regard.

## Conclusion

We conclude that it is imperative to underscore the significance of considering traditional occupation, among other pertinent factors, in the interpretation of the forthcoming substantial whole genome sequencing data. Studies have demonstrated that the farmer-related ancestry from the Iranian plateau constitutes a substantial component within the Indian gene pool. Consequently, we postulate that this Neolithic ancestry exerted an influence on the genetic composition of contemporary groups adhering to or exposed to conventional farming and pastoralist practices. We suggest that the impact of Bronze Age migration was more gradual in the Indian subcontinent than in Europe. Modern-day Indian populations include remnants of the earliest out-of-Africa migration, dispersals following the last glacial maximum, Neolithic expansion and Bronze Age migration. Therefore, a comprehensive consideration of these factors is essential when interpreting genetic data pertaining to Indian populations. The categorisation of these groups into overarching clusters such as ‘ANI’, ‘ASI’ or ‘AASI’ results in oversimplification, impeding research into rare diseases prevalent within local ancestries. Hence, we propose modeling Indian population groups based on traditional occupations, in addition to other pertinent factors.

## Limitations of the study

One limitation of this study is the use of a limited number of markers. In today’s age of whole genome sequencing technology, Y-STR analysis may be considered outdated for ancestry reconstruction. However, it is important to note that the objective of this study is not to assess individual-level variation. Instead, the study specifically assesses the impact of Neolithic agricultural expansion, which led to a significant shift in traditional occupations and the resulting endogamy in the Indian population. Our findings align with recent studies that have used whole genome data, so we believe that this limitation does not affect the overall outcome. Additionally, the non-recombinant nature and greater variability of Y-STRs provide significant statistical power for discriminating between populations.

Another challenge we encountered pertains to defining traditional occupations. This is particularly challenging because many modern populations have transitioned to urbanized lifestyles. However, a substantial portion of the population still maintains traditional practices and endogamy. This has allowed us to classify them into common clusters for the purpose of this study.

## Supporting information

ESM1

ESM2

## Acknowledgements

We are thankful to Ingenomics Private Limited for providing the Ingenomics™ Y Profiler kit.

## Author contributions

J.J.S. and G.vD. conceptualized the study. M.C., A.R.N.R., M.S., R.S., J.K., S.S., R.V.E.C. and J.J.S. performed the experiments and generated the data. J.J.S. analysed the data, performed visualisation and wrote the first draft. G.vD. edited the draft. M.S.M. and P.S. supervised the study. All the authors have reviewed and approved the final version of the submitted manuscript.

## Supplemental information

ESM1 Table S1-S2 and Figures S1–S4 EMS2 Table S1-S7

