## Supplementary material for "Y chromosome STR variation reveals traditional occupation based population structure in India": ESM1

**Footnotes:**

[1] Department of Applied Zoology, Mangalore University, Mangalagangothri, Karnataka, India
[2] Ingenomics Private Limited, New Delhi, India
[3] Institut für Sprachwissenschaft, Universität Bern, Länggassstrasse, Bern, Switzerland
[4] DNA Division, Regional Forensic Science Laboratory, Jabalpur 482011, MP, India

***Correspondence:**

* Jaison Jeevan Sequeira

** George van Driem

***Pankaj Shrivastava

### Lead Author

**Table S1.** Brief description of populations used in this study

| Population | n | Latitude | Longitude | Description |
|---|---|---|---|---|
| Kongu_Gounder | 20 | 10.89102 | 78.29115 | The traditional roles of this community are agriculture and cattle-rearing, but over time they came to be landowners, weaving, traders and money-lenders. |
| Nadar_Hindu | 31 | 8.55259 | 77.74014 | Nadar climbers were the largest subsect of today's Nadar community. A few subsects of the Nadar community, such as the Nelamaikkarars, were traditionally wealthy landlords and money lenders. Historically, most Nadars were cultivators of palmyra trees and jaggery and a few were also involved in the toddy trade. |
| Agamudayar | 18 | 9.44968 | 78.05560 | Traditionally, the Agamudayar community was involved in farming and agriculture. However, in recent times, many have diversified their occupation and have taken up businesses, jobs, and education. The community is known for its entrepreneurship and trading skills, and many are involved in buying and selling agricultural products. They are also skilled in weaving, pottery, and handicrafts. |
| Parayar | 52 | 9.81452 | 77.28149 | Malavazhiyattam is a ritualistic dance drama performed once a year by the Paraya community in Kerala. Malavazhi is the mother goddesses who are installed in the homes of the Parayas and worshiped by them. Malavazhiyattam is performed to please the deities through music and drama. |
| Tamil | 54 | 11.15506 | 78.43765 | Tamils were noted for their influence on regional trade throughout the Indian Ocean. Tamil visual art is dominated by stylized Temple architecture in major centres and the productions of images of deities in stone and bronze. |
| Haridwar | 38 | 29.95143 | 78.16947 | Mixed population |
| Rewa | 181 | 24.53978 | 81.30341 | Mixed population |
| Sarasvata_Brahmin | 179 | 30.88725 | 75.49031 | Saraswat Brahmins are brahmins who claim to have originated on the banks of river Sarasvati. As priests and religious leaders, they study religious texts, perform temple ceremonies, and conduct weddings for Hindu people of all social classes. |
| Lingayat | 97 | 14.62787 | 75.62294 | The Lingayat, also known as Devanga, are a Hindu caste from South India who traditionally work in farming, weaving, and textile merchandise. Lingayats are known for wearing small representations of a lingam, a votary object that symbolizes Shiva. |
| Vokkaliga | 102 | 12.35969 | 77.59170 | As a community of warriors and cultivators they have historically had notable |

| | | | | |
|---|---|---|---|---|
| | | | | demographic, political, and economic dominance in Old Mysore (region). |
| Himalaya | 341 | 31.23817 | 78.36184 | Mixed population |
| Himachal Pradesh | 188 | 31.73696 | 77.14238 | Mixed population |
| Odisha | 196 | 20.34546 | 84.55260 | Mixed population |
| Gujjar | 143 | 24.11184 | 73.15235 | They were traditionally involved in agriculture, pastoral and nomadic activities and formed a large heterogeneous group. |
| Uttar_Pradesh | 81 | 27.09758 | 80.92555 | Mixed population |
| Rajasthan | 309 | 26.94247 | 74.25198 | Mixed population |
| Naga | 203 | 26.12064 | 94.66182 | The Naga are tribal people from North East India. They love colour, as is evident in the shawls designed and woven by women, and in the headgear that both sexes design. Clothing patterns are traditional to each group, and the cloths are woven by the women. They use beads in variety, profusion and complexity in their jewellery. |
| Central_India | 317 | 23.23737 | 79.37815 | Mixed population |
| KT | 81 | 19.50935 | 73.78334 | Konkan tribes include Kudubi and Kharvi. They are tribal origin and reside along the Konkan coast. |
| KB | 70 | 19.35330 | 73.27925 | Konkan brahmins include Konkani speaking Sārasvata brahmins who claim to have migrated from the banks of river Sarasvati. |
| Paniya | 72 | 11.65825 | 75.82995 | The Paniya are ancient tribes of South India. The Paniya today are a scheduled tribe. One particular sub-group of theirs, the Kattupaniyar, inhabits the forest region of Nilambur in the Malappuram District. Here, members lead a traditional hunter-gatherer lifestyle. |
| Paliyan | 95 | 10.43200 | 76.81167 | They are traditional nomadic hunter-gatherers, honey hunters and foragers. Most have now transformed to traders of forest products, food cultivators and beekeepers. Some work intermittently as wage laborers, mostly on plantations. |
| Pulayar | 63 | 11.66121 | 76.43814 | Pulayars are noted for their music, craftsmanship, and for certain dances which include Kōlam-thullal, a mask dance which is part of their exorcism rituals, as well as the Mudi-āttam or hair-dance which has its origins in a fertility ritual.The folk dance Chozhikali is performed by the Pulayar community of central Kerala. |

| | | | | |
|---|---|---|---|---|
| Irula | 583 | 11.34901 5901296 927 | 78.41567 | Traditionally, the main occupation of the Irulas has been snake and rat catching, and honey collection. They also work as labourers (coolies) in the fields of the landlords during the sowing and harvesting seasons or in the rice mills. Fishing and cattle farm is also a major occupation. |
| Kadar | 28 | 10.26989 4016717 261 | 76.87759 | They are an aboriginal tribe whose traditional way of life has been based on hunting and gathering. They used to stay in the Annamalai Hills in the Western Ghats, but moved to other locations over the years. They specialized in collecting honey, wax, sago, arrowroot, cardamom, ginger, and umbrella sticks for trade with merchants from the plains. |
| Kanikaran | 17 | 10.24419 | 76.89769 | They are an aboriginal tribe whose traditional way of life has been based on hunting and gathering. They specialized in collecting honey, wax, sago, arrowroot, cardamom, ginger, and umbrella sticks for trade with merchants from the plains. |
| Thoda | 26 | 11.29005 | 78.49852 | They used to be primarily a pastoral people but now, they are increasingly venturing into agriculture and other occupations. They used to be strict vegetarians but now, some people eat meat. |
| Kota | 62 | 11.38488 | 76.76642 | They have maintained a lifestyle as jacks-of-all-trades such as potters, agriculturalist, leather workers, carpenters, and blacksmiths, and as musicians for other groups. |
| Betta Kurumba | 17 | 12.24161 | 77.45986 | They are group of Kurumbar, one of the earliest known inhabitants of the Western Ghats, who are engaged in the collection and gathering of forest produce, mainly wild honey and wax. |
| Kattunaickan | 46 | 11.66168 | 76.44050 | The Kattunayakar are one of the earliest known inhabitants of the Western Ghats, who are engaged in the hunting and gathering lifestyle. |
| Kurumba | 35 | 11.67053 | 76.42679 | The Kurumbar are one of the earliest known inhabitants of the Western Ghats, who are engaged in the collection and gathering of forest produce, mainly wild honey and wax. |
| Mullukurumba | 29 | 11.59612 | 76.26324 | They are group of Kurumbar, one of the earliest known inhabitants of the Western Ghats, who are engaged in the collection and gathering of forest produce, mainly wild honey and wax. |
| Parayar (North Tamil Nadu) | 52 | 10.23383 | 76.96315 | Paraiyars belong to the Valangai ("Right-hand caste faction"). Some of them assume the title Valangaimaan ("people of the right-hand division"). The Valangai comprised castes |

| | | | | |
|---|---|---|---|---|
| | | | | with an agricultural basis while the Idangai consisted of castes involved in manufacturing. |
| Pallar | 51 | 11.31669 | 78.25088 | There is literary evidence that suggests that Pallars were traditional farmers who produced large quantities of food grains, and that some were probably rulers in the Tamil region. |
| Paravar | 27 | 10.90704 | 76.73477 | They were maritime inhabitants of the littoral Sangam landscape known as Neithal, who were involved in pearls-harvesting, boat-building, salt-making, fishing, among other maritime activities. |
| Yadhava | 107 | 27.23442 | 77.93727 | Their traditional common function, all over India, was that of herdsmen, cowherds and milksellers. However, Jaffrelot has also said that most of the modern Yadavs are cultivators, mainly engaged in tilling the land, and less than one third of the population are occupied in raising cattle or the milk business. |
| Vanniyar | 21 | 10.72359 | 77.66860 | Most Vanniyars remain either marginal farmers cultivating small areas of land or landless labourers. |
| Vanniyar (North Tamilandu) | 96 | 10.78835 | 77.71255 | Most Vanniyars remain either marginal farmers cultivating small areas of land or landless labourers. |
| Nadar (Tiruneveli) | 59 | 8.625215 | 77.87851 | A few subsects of the Nadar community, such as the Nelamaikkarars, were traditionally wealthy landlords and money lenders. Historically, most Nadars were cultivators of palmyra trees and jaggery and a few were also involved in the toddy trade. |
| Cape Nadar | 98 | 8.66866 | 77.74714 | Their traditional occupation was climbing trees and gathering the sap of coconuts to make palm wine. |
| Piramalai Kallar | 53 | 11.12272 | 78.55850 | By the late 18th century, the Kallars were working as kavalkarars, or watchmen, in hundreds of villages throughout southern Tamil Nadu, especially the region west of Madurai. These kavalkarars were given maniyam, rent-free land, to ensure they did their job correctly. |
| Maravar | 80 | 11.38132 | 78.56948 | The Maravar community, along with the Kallars, had a reputation for thieving and robbery from as early as the medieval period. |
| Valayar | 95 | 11.28437 | 78.54751 | They are traditionally a community of net weavers. |
| Tamil Jains | 100 | 11.18739 | 78.51455 | The traditional occupation of the majority of the Tamil Jain families has been landowners of agricultural land. |
| Ezhava | 95 | 10.84639 | 76.46202 | The Ezhava used to work as agricultural labourers, small cultivators, toddy tappers; some were also involved in weaving and some practised Ayurveda. |
| Mukkuvar | 17 | 10.80323 | 76.83556 | Their traditional occupation is diving for pearls and seashell. |

| | | | | |
|---|---|---|---|---|
| Sourashtra | 40 | 12.75646 | 78.23989 | They are known for their expertise in traditional vedic practices and rituals. Apart from being priests and scholars, they are also been involved in various professions such as agriculture, trade, and business and were a prominent industrious and prosperous mercantile community of merchants and weavers in southern India until the 20th century. |
| Brahacharanam Brahmin | 21 | 11.12272 | 78.55850 | The traditional function of Brahacharanams is to study and impart Vedic knowledge or officiate as priests in religious functions. |
| Iyengar Brahmin | 11 | 12.76691 | 78.22860 | Iyengars are an ethnoreligious community of Tamil-speaking Hindu Brahmins, whose members follow Sri Vaishnavism and the Visishtadvaita philosophy propounded by Ramanuja. |
| Vadama Brahmin | 63 | 13.01852 | 78.70048 | The term "vadama" may refer to proficiency in Sanskrit and Vedic rituals. Sociologist Andre Beteille in his thesis ‘Caste, class, and power: changing patterns of stratification in a Tanjore village’, describes them as the biggest mirasidars (landlords) among the Iyer community. |
| Bhotra | 133 | 21.29021 | 83.24523 | Almost all Bhotra/Bhottada live in the Indian state of Odisha. They are thought to be among the original inhabitants of this region of India. In the distant past the Bhottada were a hunter-gatherer people. |
| Haryana Brahmin | 99 | 28.68738 | 75.55483 | These are brahmins from a northern Indian state named Haryana. |

**Table S2.** Table showing frequency of major haplogroups found in the studied populations based on YSTR data

| Groups | Haplogroups (%) | | | | | | | | | | | | Cluster |
|---|---|---|---|---|---|---|---|---|---|---|---|---|---|
| | **C** | **E1** | **F** | **G** | **H** | **I** | **J2** | **L** | **Q** | **R1a** | **R1b** | **R2** | |
| Dravidian | 3.2 | 2.4 | 10.8 | 3.0 | 22.3 | 1.6 | 10.0 | 17.0 | 1.4 | 13.4 | 0.0 | 9.8 | Linguistic |
| Indo-European | 0.63 | 7.64 | 0.0 | 2 | 16.66 | 2.82 | 9.2 | 10.39 | 5.01 | 31.75 | 0.5 | 3.26 | Linguistic |
| Priestly | 1.2 | 2.9 | 0.2 | 2.4 | 10.7 | 1.9 | 10.0 | 10.2 | 6.6 | 43.4 | 0.7 | 4.3 | Occupation |
| Farming-related | 2.6 | 3.3 | 5.5 | 4.8 | 19.5 | 0.5 | 9.7 | 25.6 | 1.8 | 13.0 | 0.0 | 10.6 | Occupation |
| Hunter-gatherer | 2.6 | 3.9 | 37.7 | 1.4 | 21.3 | 2.8 | 6.9 | 15.9 | 3.6 | 1.5 | 2.6 | 6.2 | Occupation |
| Tribes | 2.6 | 3.9 | 14.7 | 1.4 | 21.3 | 2.8 | 6.9 | 15.9 | 3.6 | 10.7 | 2.6 | 6.2 | Social |
| Castes | 2.2 | 3.0 | 2.8 | 3.6 | 17.7 | 1.2 | 10.9 | 19.2 | 3.1 | 23.3 | 0.3 | 8.7 | Social |
| Iran-West | 1.21 | 10.92 | 0.00 | 8.84 | 2.60 | 1.04 | 33.97 | 6.07 | 8.32 | 12.48 | 7.63 | 2.77 | Iran |
| Iran-East | 0.0 | 0.0 | 0.0 | 0.8 | 0.0 | 2.7 | 0.8 | 52 | 5.5 | 4.3 | 3.9 | 0.0 | Iran |
| Europe | 0.0 | 6.25 | 3.13 | 0.52 | 0.00 | 25.52 | 5.21 | 0.52 | 2.08 | 43.23 | 7.29 | 0.0 | Europe |

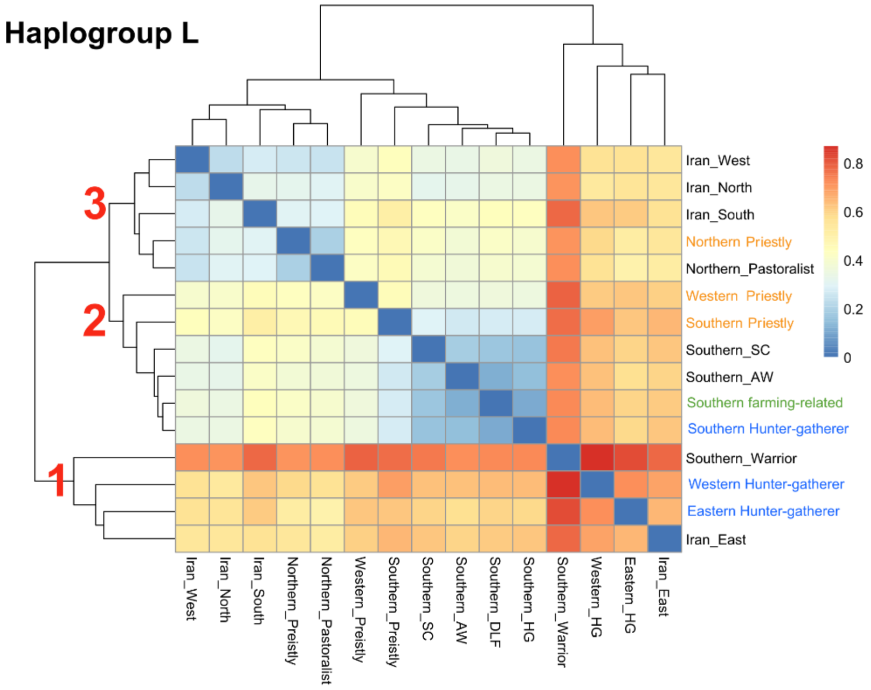


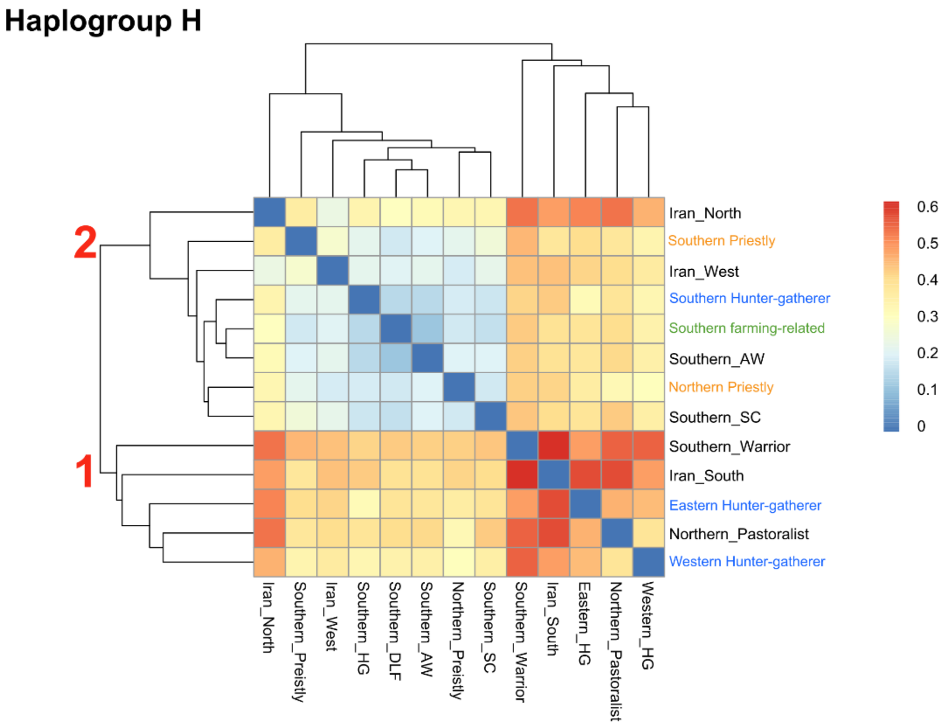


**Figure S1**. RST based Heatmap for H and L samples. Three clusters are shown in the figure. Blue represents hunter-gatherer group, orange represents priestly group and green represents farming-related group.

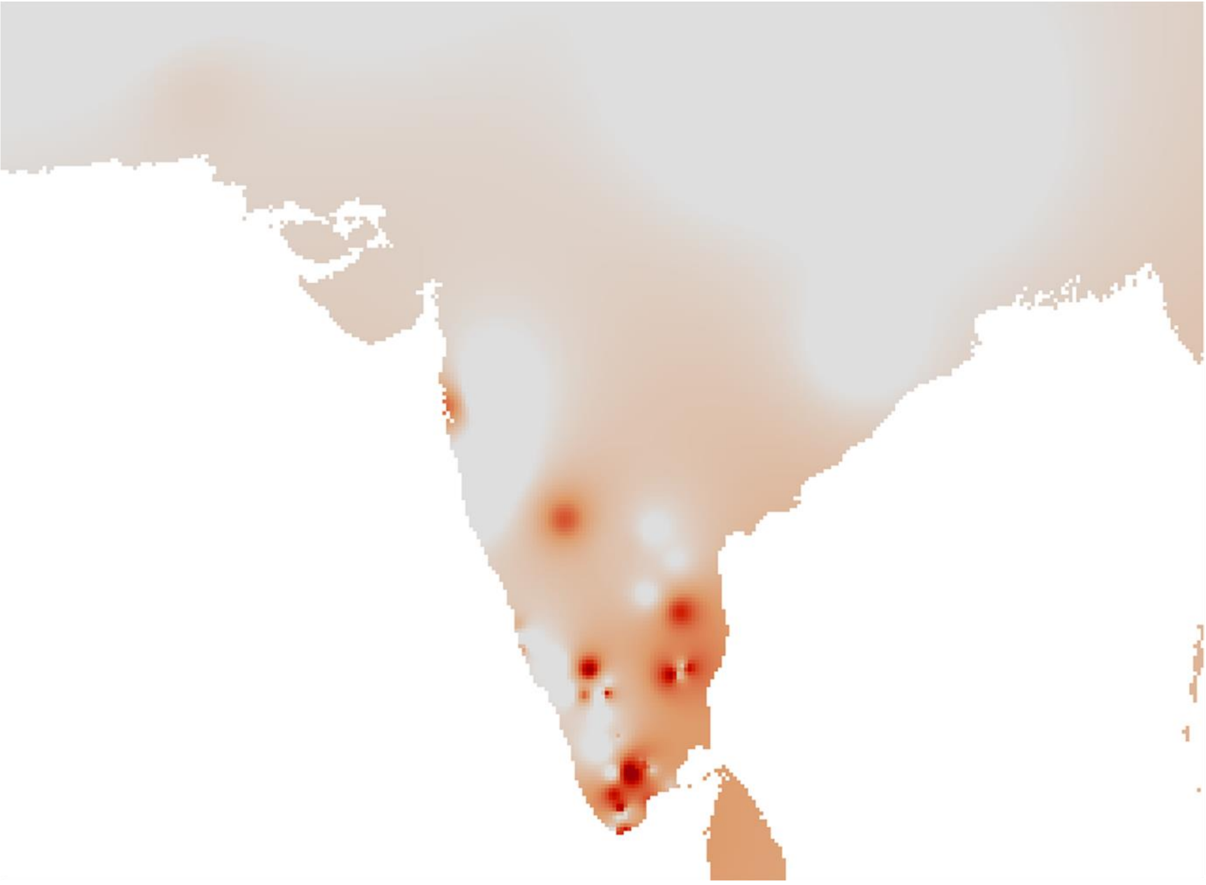

**Figure S2.** Spatial distribution of L-M27/M76 haplogroup in India

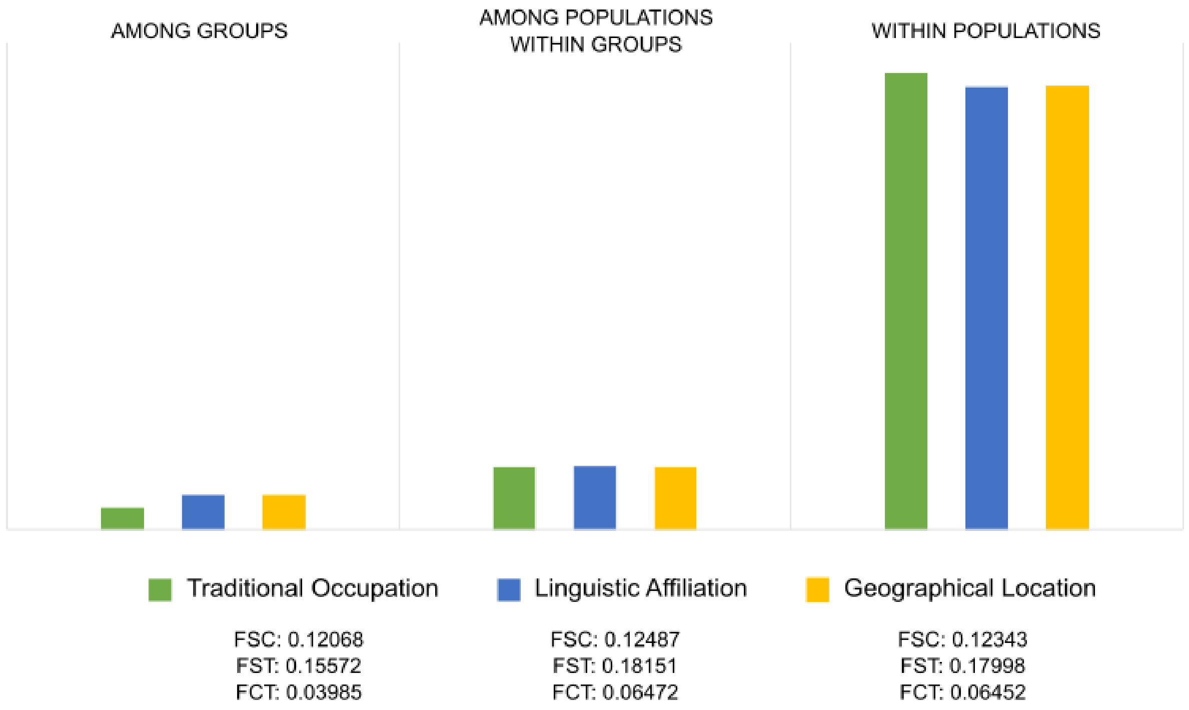


**Figure S3**. AMOVA results showing variance contributed by linguistic differences, geographical distance and traditional occupation.

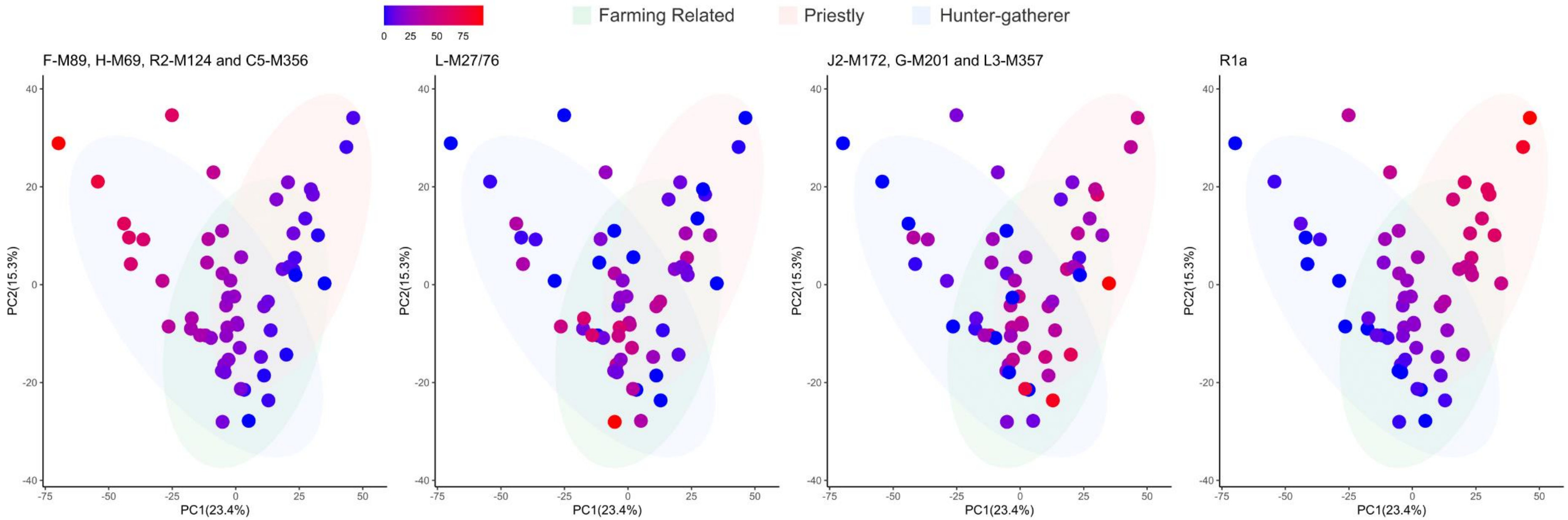


**Figure S4.** Gradient of haplogroup frequency showing the contribution of different haplogroups in the PCA plot.
